# Finding drug targeting mechanisms with genetic evidence for Parkinson’s disease

**DOI:** 10.1101/2020.07.24.208975

**Authors:** Catherine S. Storm, Demis A. Kia, Mona Almramhi, Sara Bandres-Ciga, Chris Finan, Aroon D. Hingorani, International Parkinson’s Disease Genomics Consortium (IPDGC), Nicholas W. Wood

## Abstract

Parkinson’s disease (PD) is a neurodegenerative movement disorder that currently has no disease-modifying treatment, partly owing to inefficiencies in drug target identification and validation using human evidence. Here, we use Mendelian randomization to investigate more than 3000 genes that encode druggable proteins, seeking to predict their efficacy as drug targets for PD. We use expression and protein quantitative trait loci for druggable genes to mimic exposure to medications, and we examine the causal effect on PD risk (in two large case-control cohorts), PD age at onset and progression. We propose 23 potential drug targeting mechanisms for PD, of which four are repurposing opportunities of already-licensed or clinical-phase drugs. We identify two drugs which may increase PD risk. Importantly, there is remarkably little overlap between our MR-supported drug targeting mechanisms to prevent PD and those that reduce PD progression, suggesting that molecular mechanisms driving disease risk and progression differ. Drugs with genetic support are considerably more likely to be successful in clinical trials, and we provide compelling genetic evidence and an analysis pipeline that can be used to prioritise drug development efforts for PD.

## Introduction

Parkinson’s disease (PD) is a neurodegenerative movement disorder that currently has no disease-modifying treatment. Despite efforts, around 90% of drugs that enter clinical trials fail, mostly due to insufficient efficacy or safety (Wouters, McKee, and Luyten 2020; Smietana, Siatkowski, and Møller 2016; Harrison 2016). This contributes to the staggering $1.3 billion mean price of bringing a new drug to the market (Wouters, McKee, and Luyten 2020).

Incorporating genetics in drug development could be one of the most efficient ways to improve this process, since drugs with genetic support are considerably more likely to succeed in clinical trials (Nelson et al. 2015; King, Wade Davis, and Degner 2019; Hingorani et al. 2019). So-called “druggable” genes encode proteins that can be targeted by medications. More precisely, these are proteins which have been targeted by a pharmacological agent or are considered possible to target with a small molecule or monoclonal antibody (Finan et al. 2017; Schmidt et al. 2020). While genome-wide association studies (GWAS) have effectively identified single nucleotide polymorphisms (SNPs) linked to PD risk and progression (Nalls et al. 2019; Blauwendraat et al. 2019; Iwaki et al. 2019), the GWAS design cannot reliably pinpoint causal genes nor directly inform drug development.

Mendelian randomization (MR) is a genetic technique that can predict the efficacy of a drug by mimicking a randomized controlled trial (Katan 1986; Smith and Ebrahim 2003; Hingorani et al. 2005; Holmes et al. 2017). MR can use genetic variants, usually SNPs, associated with expression levels of a druggable gene to mimic lifelong exposure to a medication targeting the encoded protein (Schmidt et al. 2020; Zhu et al. 2016). The association between the same genetic variants and a disease (the outcome) can then be extracted from a GWAS (Figure 1a). The SNP-gene and SNP-disease associations can be combined to infer the causal effect of the exposure on the outcome. The exposure and outcome can be measured in two independent cohorts, meaning that openly available data from two large-scale GWASs can be used for one well-powered MR study. Because of Mendel’s law of independent assortment, individuals are “randomized” at conception to have genetically higher or lower expression levels of the druggable gene (Figure 1b). Individuals are generally unaware of their genotype, so the MR study is effectively blinded.

**Figure 1:**
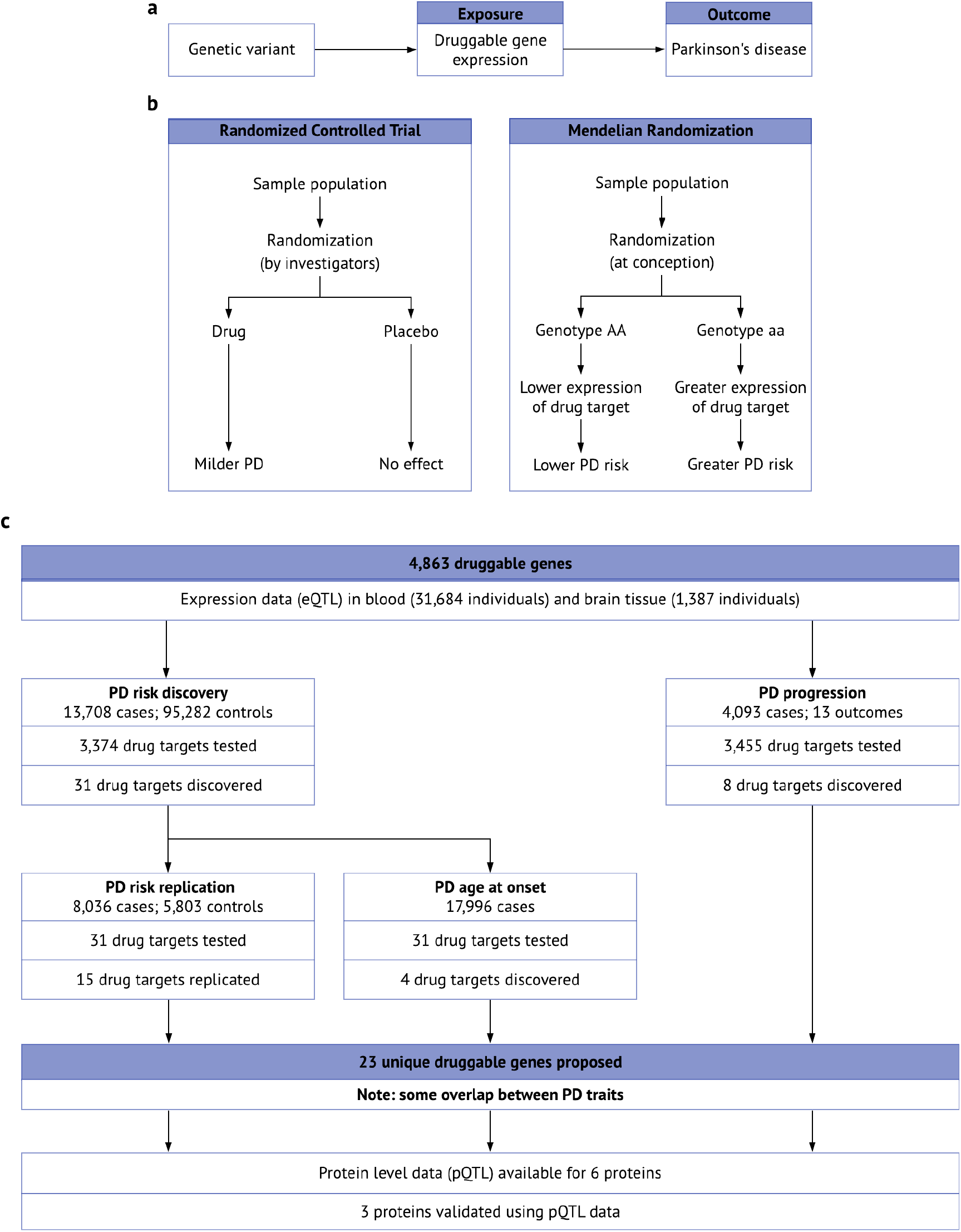
Overview of MR and our study. a. Genetic variants associated with the expression of a gene are called eQTLs, and they mimic life-long exposure to higher or lower levels of gene expression (the exposure). These variants affect PD (the outcome) through the exposure only, i.e. there is no horizontal pleiotropy. b. MR is analogous to a randomized controlled trial, where individuals are randomly allocated to a genotype according to Mendel’s law of independent assortment. Adapted from (Hingorani et al. 2005). c. Workflow and summarized results of our MR study.

As the MR literature grows and more robust methods arise, the potential of MR in drug development for neurodegenerative disease has become increasingly clear (Burgess et al. 2019; Storm et al. 2020). At the same time, large-scale GWASs for PD risk and progression markers have become available (Nalls et al. 2019; Blauwendraat et al. 2019; Iwaki et al. 2019). For the first time, it is possible to combine these resources to identify drug targets for PD with genetic support.

In this study, we predict the efficacy of over 3000 drug targeting mechanisms in PD using gene expression in blood and brain tissue to mimic the action of medications. Figure 1c gives a schematic overview of our analysis. We investigate the causal effect of gene expression on PD risk in two independent case-control cohorts and on a range of PD progression markers. Where possible, we repeat the analysis using SNPs associated with circulating levels of the encoded proteins. Using large-scale, openly available GWAS data and modern Mendelian randomization techniques, we propose a list of genetically-supported drug targets for PD, including repurposing opportunities of already-licensed or clinical-phase drugs.

## Results

### Mimicking medications with expression quantitative trait loci

Conceptually, the druggable genome encompasses all human genes that encode drug targets, and the most comprehensive version of the druggable genome to date includes 4,863 genes. These genes code for druggable proteins, including proteins targeted by approved and clinical-phase drugs, proteins similar to approved drug targets as well as proteins accessible to monoclonal antibodies or drug-like small molecules *in vivo* (Finan et al. 2017).

An expression quantitative trait locus (eQTL) is a genetic variant associated with expression levels of a gene (Figure 1a). We sought to identify openly available eQTL data for 4,863 druggable genes to mimic exposure to the corresponding medications (Finan et al. 2017). For example, an eQTL associated with reduced expression of *HMGCR* mimics exposure to an HMGCR-inhibitor, such as a statin. Most clinically-used drugs target proteins, and genetic variants associated with protein levels in a clinically relevant tissue may be ideal to model drug target effects with MR (Schmidt et al 2020). However, even with high throughput protein assays, the spectrum of reliable, well-powered GWAS data on protein targets is limited. Additionally, many genetic studies on protein levels are based on plasma and lack any tissue specificity (Sun et al. 2018; Emilsson et al. 2018; Suhre et al. 2017). Therefore, gene expression levels can be used to proxy the drug target. Although the transcript level is biologically one step before the protein level, expression GWAS studies cover many more genes across the genome and provide tissue specificity. As such, using gene expression data provides a very good resource for high level screens to develop drug targeting hypotheses.

We used eQTL data from the eQTLGen consortium to imitate drug action in blood, which include 31,684 mostly European-ancestry individuals (Võsa et al. 2018). We also used brain tissue eQTL data provided by the PsychENCODE consortium, which are based on 1,387 prefrontal cortex samples of mostly European ancestry (679 healthy controls, 497 schizophrenia, 172 bipolar disorder, 31 autism spectrum disorder, 8 affective disorder patients) (Wang et al. 2018). We only kept eQTLs with false discovery rate (FDR) < 0.05 and located within 5 kb of the associated gene to maximise the specificity of the eQTL.

Overall, eQTLs within 5 kb of the gene were available for 2,786 and 2,448 druggable genes in blood and brain tissue, respectively. These were clumped at *r*^2^ = 0.2, and a linkage disequilibrium matrix based on the 1000 genomes EUR reference panel was included in the MR analysis to account for correlation between genetic variants (Burgess, Dudbridge, and Thompson 2016; The 1000 Genomes Project Consortium 2012).

### Discovery phase identifies 31 potential drug targets to prevent PD

The largest GWAS dataset available for a PD trait is disease risk in individuals of European descent, obtained from the International PD Genomics Consortium (IPDGC) (Nalls et al. 2019). Our discovery cohort consisted of samples collected for a 2014 GWAS meta-analysis, including 13,708 PD patients and 95,282 controls (Nalls et al. 2014). In this dataset, eQTLs for 2,689 and 2,256 genes were available for MR in blood and brain tissue, respectively. The MR effect estimate for each SNP (Wald ratio) was calculated, and where > 1 eQTL was available per gene (after clumping at *r*^2^ = 0.2), Wald ratios were meta-analysed, weighted by inverse-variance (IVW). Expression of 14 genes in blood and 20 in brain tissue was significantly associated with PD risk at FDR < 0.05 (Table S1). All of these remained significant when clumping at *r*^2^ = 0.001 (Table S1). Overall, genetically-determined expression of 31 genes (11 in blood only; 17 in brain tissue only; 3 in both blood and brain tissue) was significantly associated with PD risk in the discovery cohort at FDR < 0.05.

### 15 potential preventative agents replicate in an independent PD case-control cohort

Replication between independent cohorts has been essential in genetics to establish the credibility of genotype-phenotype associations (Hirshhorn et al. 2002; Chanock et al. 2007; Marigorta et al. 2018). Although this lesson has been of utmost importance, very few MR studies to date attempt replication (Burgess, Foley, and Zuber 2018), perhaps because several independent GWAS datasets for the same trait are often not readily available. We investigated all genes which reached significance in the discovery phase using the Wald ratio or IVW method in an independent PD case-control cohort (Figure 1). The replication population consisted of 8,036 PD patients and 5,803 controls, with no overlap with the discovery cohort (Nalls et al. 2019). The MR methods were identical to those used in the discovery phase.

Genetically-predicted expression of 15 genes (4 in blood only; 9 in brain tissue only; 2 in both tissues) replicated using the Wald ratio or IVW method (Figure 2, Table S1). *BST1, CD38, CHRNB1, CTSB, GPNMB, HSD3B7, LDADLS3, MAPT, MMRN1, NDUFAF2, PIGF, VKORC1* and *WNT3* reached FDR < 0.05, and *ACVR2A* and *MAP3K12* reached nominal significance. *GPNMB* and *HSD3B7* reached significance in both blood and brain tissue. Of these 15 potential drug targets to prevent PD, 10 were not nominated by the PD risk GWAS meta-analysis (Nalls et al. 2019), illustrating the added value of this MR approach.

**Figure 2:**
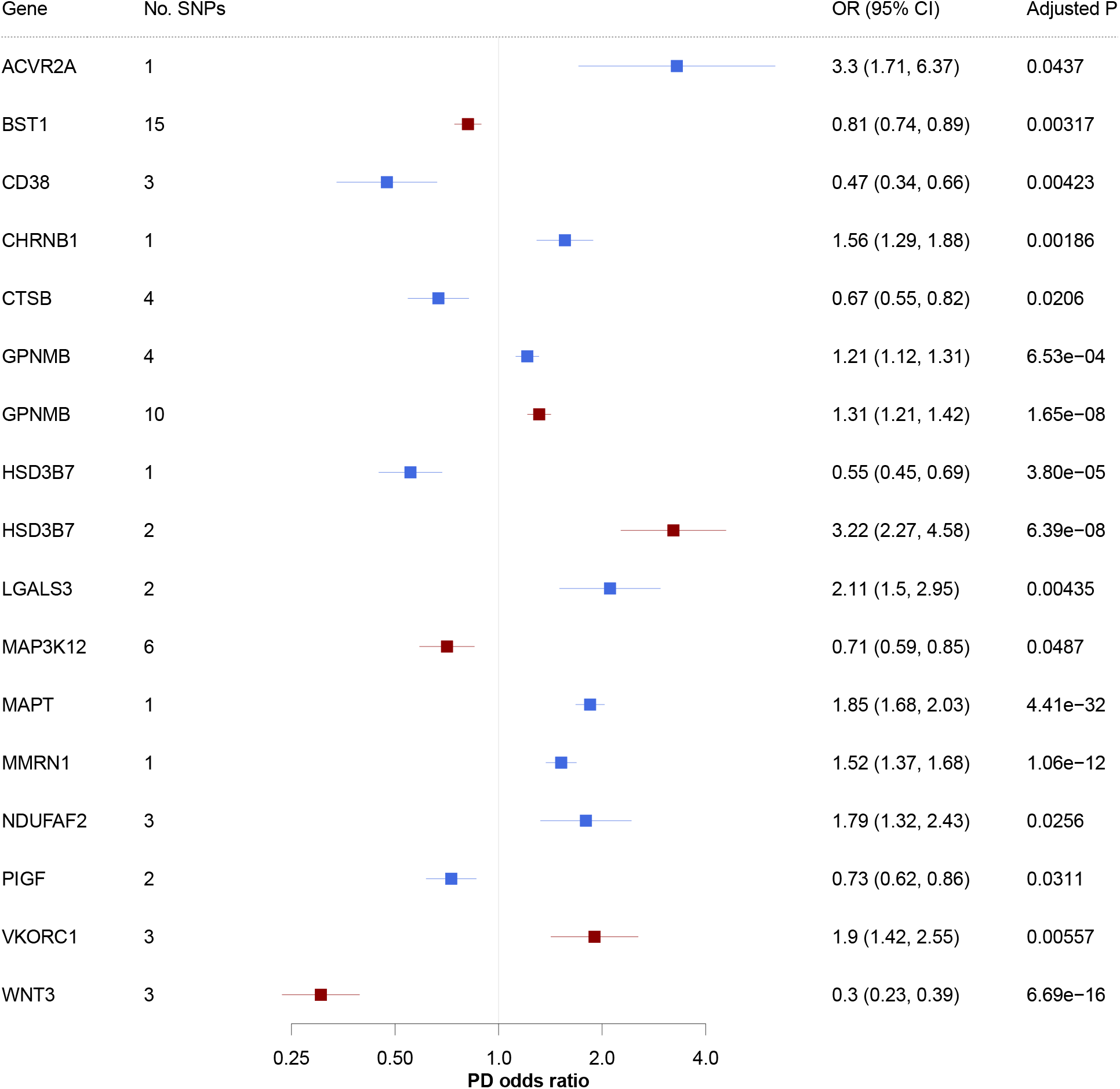
15 potential preventative drug targets reach significance in two independent PD case-control cohorts. Forest plots showing the discovery-phase results for the 16 replicated genes. PD odds ratio per 1-standard-deviation increase in gene expression. Results from the Wald ratio or IVW are shown and colour-coded according to the tissue (red = blood, blue = brain tissue). 95% CI, 95% confidence interval; OR, odds ratio.

Six replicated genes encode targets of approved or clinical-phase drugs. Three of these are targeted by a drug with an appropriate direction of effect for PD protection, *CHRNB1*, *NDUFAF2* and *VKORC1* (Table 1 and S1), meaning that there may be a repurposing opportunity for the corresponding drugs. The GPNMB protein is a receptor targeted by glematumumab, an antibody-drug conjugate that is being evaluated for several types of cancer (Rose et al. 2017). After binding to GPNMB, the drug is internalised by the cell and is cytotoxic. Since this mechanism of action does not reflect a change in GPNMB levels, we do not consider glematumumab a potential candidate for repurposing.

**Table 1:**
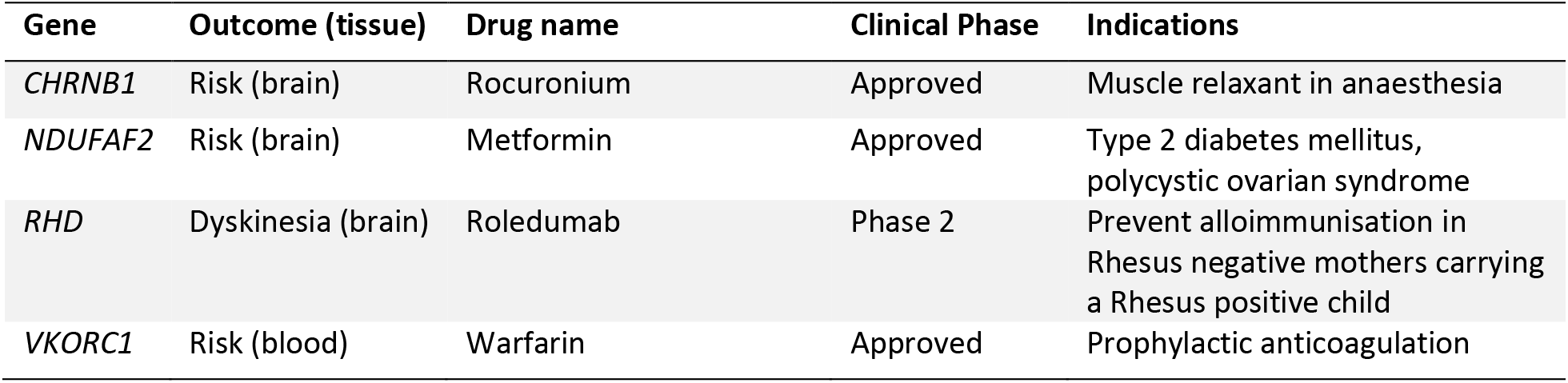
Four potential drug targeting mechanisms for PD may constitute repurposing opportunities for existing drugs. These drugs are either approved or in clinical trial phase, and the mechanism of action is consistent with direction of our MR effect estimate. The second column displays the potential effect on PD and target tissue. Clinical phase and drug indication based on https://clinicaltrials.gov/ and the British National Formulary. Direction of effect was confirmed using https://www.drugbank.ca or https://www.ebi.ac.uk/chembl/databases.

We find that CD38-inhibitors such as daratumumab, licensed to treat multiple myeloma, may increase PD risk. We also find that MAP3K12-inhibitors such as CEP-1347 may increase PD risk. Interestingly, CEP-1347 was investigated as a PD treatment, and our data provide an explanation why this drug was not effective (Parkinson Study Group PRECEPT Investigators 2008). As such, this MR approach identifies not only potential drug targets and repurposing opportunities, but also licensed medications which may raise disease risk.

### MR quality control suggests that *CD38, GPNMB*, and *MAP3K12* have the most robust MR evidence for PD risk

We completed a series of quality control steps to prioritise the replicated genes. Firstly, the direction of effect was consistent between the discovery and replication phases for all 15 replicated genes (Table S2). We note that genetically-predicted expression of *HSD3B7* was associated with raised PD risk in blood, but reduced PD risk in brain tissue (Figure 2). This pattern occurred both in the discovery and replication phase. Although this may suggest opposing effects between tissues, there was only one eQTL available for *HSD3B7* in brain and two eQTLs in blood (discovery phase). It is therefore not possible to perform the additional quality control discussed below, which illustrates that results based on one or two SNPs should be interpreted with caution. It is noteworthy that previous eQTL-based MR studies have reported heterogeneity between tissues, both in terms of the magnitude and direction of effect (Schmidt et al. 2020)

We consider the MR result more robust if several meta-analysis methods yield a similar result, such as the maximum likelihood and MR-Egger methods (Haycock et al. 2016; Burgess et al. 2019; Slob and Burgess 2020). This is only possible if >2 SNPs are available per gene, and we found that all eight genes with > 2 SNPs reached at least nominal significance using the maximum likelihood method (unadjusted *p* < 0.05). The magnitude and direction of effect was largely consistent between methods, except for *BST1*. For *BST1*, the MR-Egger estimate was in the opposite direction to the IVW and maximum likelihood results (Table S1), and we further discuss our interpretation of this later in the text.

MR assumes that the SNP only affects the outcome (PD risk) through the exposure (gene expression), and therefore the y-intercept of the IVW regression is fixed at zero (Burgess et al. 2019). This assumption is violated if there is genetic pleiotropy, where a SNP affects the outcome through an alternative pathway. If genetic pleiotropy pushes the effect in one direction, the IVW method will be biased. The MR-Egger method relaxes this assumption by not constraining the y-intercept. If the MR-Egger y-intercept significantly deviates from zero, it suggests that there is directional pleiotropy. Of the eight genes with > 2 SNPs available, all passed the MR-Egger intercept test except *BST1*, explaining the deviant MR-Egger estimate for this gene (Table S2).

Nevertheless, if SNPs for the same gene are pleiotropic in opposing directions, the MR-Egger y-intercept will still be zero. Here, the Cochran’s Q and *I*^2^ tests are useful, which assess overall heterogeneity between Wald ratios. Here, *NDUFAF2, WNT3* and *VKORC1* did not pass the Cochran’s Q test (*p* < 0.05), and six genes did not pass the *I*^2^ test (*I*^2^ > 0.50): *BST1, CTSB, GPNMB* (in brain tissue), *NDUFAF2, VKORC1*, and *WNT3* (Table S2).

Additionally, an MR result may be driven by a locus where the SNP-exposure and SNP-outcome associations are rooted in two distinct causal SNPs (Hemani et al. 2018a). In other words, the SNP driving the exposure may be in close linkage disequilibrium with the SNP driving the outcome. This can be probed using colocalization analysis (Giambartolomei et al. 2014). Using a colocalization approach, Kia and colleagues recently found that eQTLs in brain tissue for *CD38* and *GPNMB* colocalize with PD risk loci (Kia et al. 2020). Notably, the eQTL datasets used by the authors (Ramasamy et al. 2014, The GTEx Consortium 2015) differ from those in this study (Wang et al. 2018, Võsa et al. 2018). This colocalization evidence strengthens the evidence for CD38 and GPNMB as drug targets for PD.

Overall, three genes had consistent effects between cohorts and meta-analysis methods, and they passed the MR-Egger intercept, Cochran’s Q and *I*^2^ tests in the discovery phase. As such, these genes carry the most robust MR evidence for a causal relationship with PD risk: *CD38, GPNMB* and *MAP3K12*.

### Four potential targets for preventative drugs may also affect PD age at onset

Pharmacologically delaying the age of onset of a debilitating disease may have a considerable impact on the quality of affected individuals’ lives, providing disability-free years to people at risk. Evidence from polygenic risk score analyses suggest that genetic risk of PD is correlated with PD age at onset (Escott-Price et al. 2015; Nalls et al. 2015; Ibanez et al. 2017; Blauwendraat et al. 2019). We therefore asked whether expression of the genes reaching significance in our MR discovery phase for PD risk also predict PD age of onset. We performed the MR analysis for these genes using openly available summary statistics from a PD age of onset GWAS, including 17,996 PD patients (Figure 1). Based on the same analysis pipeline as the replication step for PD risk, we found that expression of four genes predicted PD age of onset at *p* < 0.05: *BST1* in blood*, CD38* in brain tissue, *CTSB* in brain tissue and *MMRN1* in brain tissue (Figure 3, Table S3). *CD38* and *MMRN1* remained significant when clumping at *r*^2^ = 0.001. There were > 2 SNPs available *for BST1, CD38* and *CTSB*, and the IVW, maximum likelihood and MR-Egger methods yielded a consistent direction of effect (Table S1). All three genes passed the MR-Egger intercept (*p* > 0.05), and Cochran’s Q test (*p* > 0.05), whereas none passed the *I*^2^ test (*I*^2^ > 0.50).

**Figure 3:**
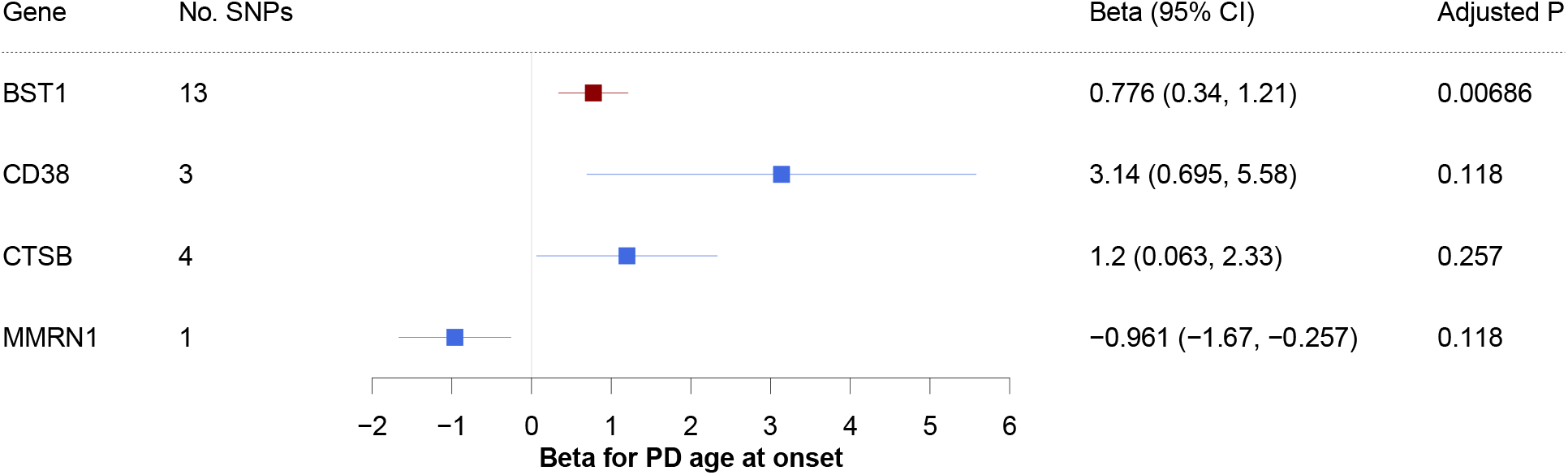
Four potential preventative drugs may also affect PD age at onset. Forest plot showing standard-deviation change in PD age at onset per 1-standard-deviation increase in gene expression. A negative beta corresponds to a younger age at onset, and a positive beta corresponds to an older age at onset. Results from the Wald ratio or IVW method are shown, colour-coded by tissue (red = blood, blue = brain tissue). 95% CI, 95% confidence interval.

We hypothesized that if increased expression of a gene predicts reduced PD risk, this gene should be associated with a delayed age at onset. This was consistently the case for all four genes that reached significance for age at onset. Overall, these data suggest that there may be some shared molecular mechanisms driving PD risk and age at onset, and that this overlap is incomplete.

### There is little overlap between drug targets to prevent PD and reduce PD progression

The PD risk GWAS data afford large discovery and replication cohorts, which is a great advantage in MR. Nevertheless, it is currently not possible to reliably predict PD, limiting the immediate usefulness of a drug to prevent or delay this condition. Many clinical trials for PD use progression markers such as the Unified PD Rating Scale (UPDRS) to evaluate efficacy of a drug, and it remains unclear how the molecular mechanisms driving PD risk relate to clinical progression. We therefore used MR to probe whether expression of any of the 4,863 druggable genes is significantly associated with PD progression, measured by the UPDRS (total and parts 1 to 4), mini-mental state examination (MMSE), Montreal cognitive assessment (MOCA), modified Schwab and England activities of daily living scale (SEADL), Hoehn and Yahr stage, dementia, depression, and dyskinesia. The MR pipeline for each progression marker was identical to the discovery phase for PD risk (Figure 1).

We used openly available summary statistics from a GWAS for these PD progression markers, which included 4,093 European PD patients, followed over a median of 2.97 years (Iwaki et al. 2019). 3,455 genes had eQTLs in blood available for MR analysis using any of the 13 PD progression markers (2,752 in blood, 2,353 in brain tissue), and eight genes reached significance across five progression outcomes (Figure 4, Table S1). Of these, one gene, *RHD*, encodes the target of a clinical-phase medication with an appropriate direction of effect, providing a potential repurposing opportunity (Table 1).

**Figure 4:**
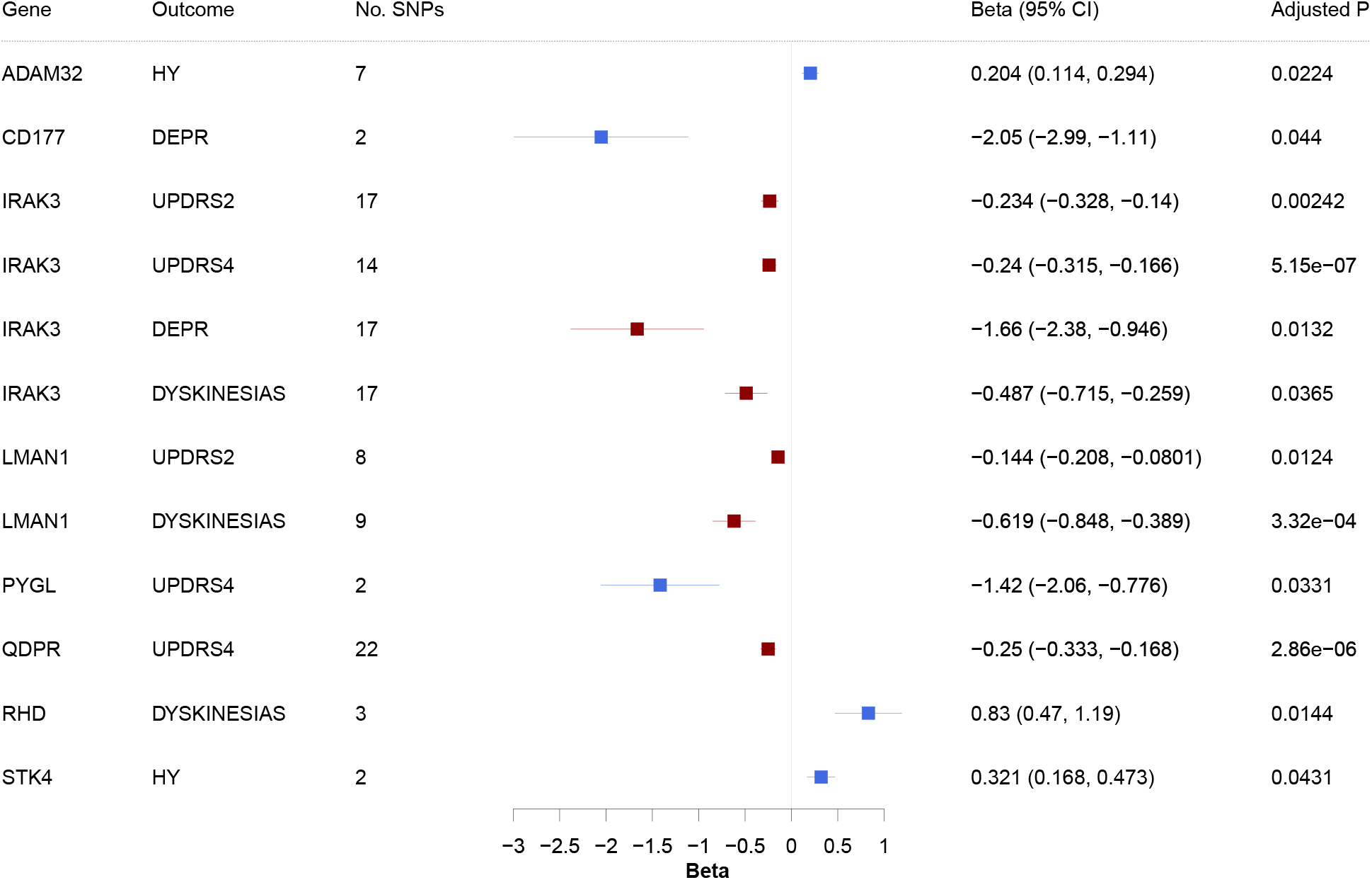
Genetically-predicted expression of eight genes in blood or brain tissue is associated with PD progression markers. Forest plot showing the standard-deviation change in each progression marker, per 1-standard-deviation increase in gene expression. Results from the Wald ratio or IVW method are shown and colour-coded according to the tissue (red = blood, blue = brain tissue). HY, Hoehn and Yahr; DEPR, depression, UPDRS2-4, unified PD rating scale parts 2 and 4.

No genes reached significance in both brain and blood tissue. Genetically-predicted *IRAK3* expression in blood was significantly associated with four different progression outcomes (UPDRS parts 2 and 4, depression, and dyskinesias), and *LMAN1*-expression in blood reached significance for both dyskinesias and UPDRS part 2. This strengthens the evidence for these two genes.

We performed the same quality control as in the PD risk analysis. The direction of effect was consistent between the IVW, maximum likelihood and MR-Egger methods for all genes except *RHD*, where the MR-Egger method opposed the direction of the IVW and maximum likelihood methods. In brain tissue, *CD177* (depression), *RHD* (dyskinesias), *PYGL* (UPDRS part 4) and *STK4* (Hoehn and Yahr) reached significance when clumping at *r*^2^ = 0.001. The MR-Egger intercept, Cochran’s Q and *I*^2^ tests were possible for 10 gene-outcome combinations. The genes *IRAK3* (dyskinesia), *LMAN1* (UPDRS part 2), *QDPR* (UPDRS part 4) and *RHD* (dyskinesia) passed all three of these quality control tests for pleiotropy (Table S2). No genes reached significance for both PD risk and these progression markers. Taken together, the genes *IRAK3, LMAN1, QDPR* and *RHD* have the most robust MR evidence for modifying a marker of PD progression.

### Protein quantitative trait locus data provide further genetic evidence

Expression QTL data provide an ideal resource for high-level screens to develop drug targeting hypotheses, providing data for a great number of genes across many tissues (Võsa et al. 2018; Wang et al. 2018). Nevertheless, most clinically-used drugs target proteins, not gene expression. As such, genetic variants associated with protein levels, called protein quantitative trait loci (pQTLs), may model drug target effects more accurately (Schmidt et al 2020). We therefore sought to validate our 23 proposed drug targets for PD using pQTL data. Well-powered and tissue diverse pQTL data are however limited; for PD risk, we found pQTLs for four of our proteins of interest: BST1, CTSB, GPNMB, LGALS3 (Sun et al. 2018; Emilsson et al. 2018; Suhre et al. 2017). For the progression outcomes, we found pQTLs for PYGL and QDPR, which our eQTL study identified as drug target candidates using UPDRS part 4 as the outcome (Sun et al. 2018; Emilsson et al. 2018).

For these six proteins with available pQTLs, our MR analysis found that three were associated with PD risk or UPDRS part 4 at nominal significance (unadjusted *p* < 0.05): BST1, CTSB and LGALS3 (Figure 5, Table S5). Importantly, we find similar results for these proteins when using data from different pQTL studies. In contrast, the result was not consistently significant for GPNMB and PYGL when using pQTLs identified by different pQTL studies. For PYGL, the two pQTLs discovered by different pQTL studies, rs62143198 and rs2297890, are located on chromosomes 19 and 14, respectively (Sun et al. 2018; Emilsson et al. 2018). The PYGL gene is located on chromosome 14. The pQTL on chromosome 14 reached nominal significance, but this effect was not in the same direction as the eQTL result in brain tissue (Figure 4). Whereas the pQTL on chromosome 19 tended in the direction of the eQTL result, this result did not reach significance. The small number of pQTLs available per protein limits the possibility to perform MR quality control, making is difficult to evaluate the robustness of this result.

**Figure 5:**
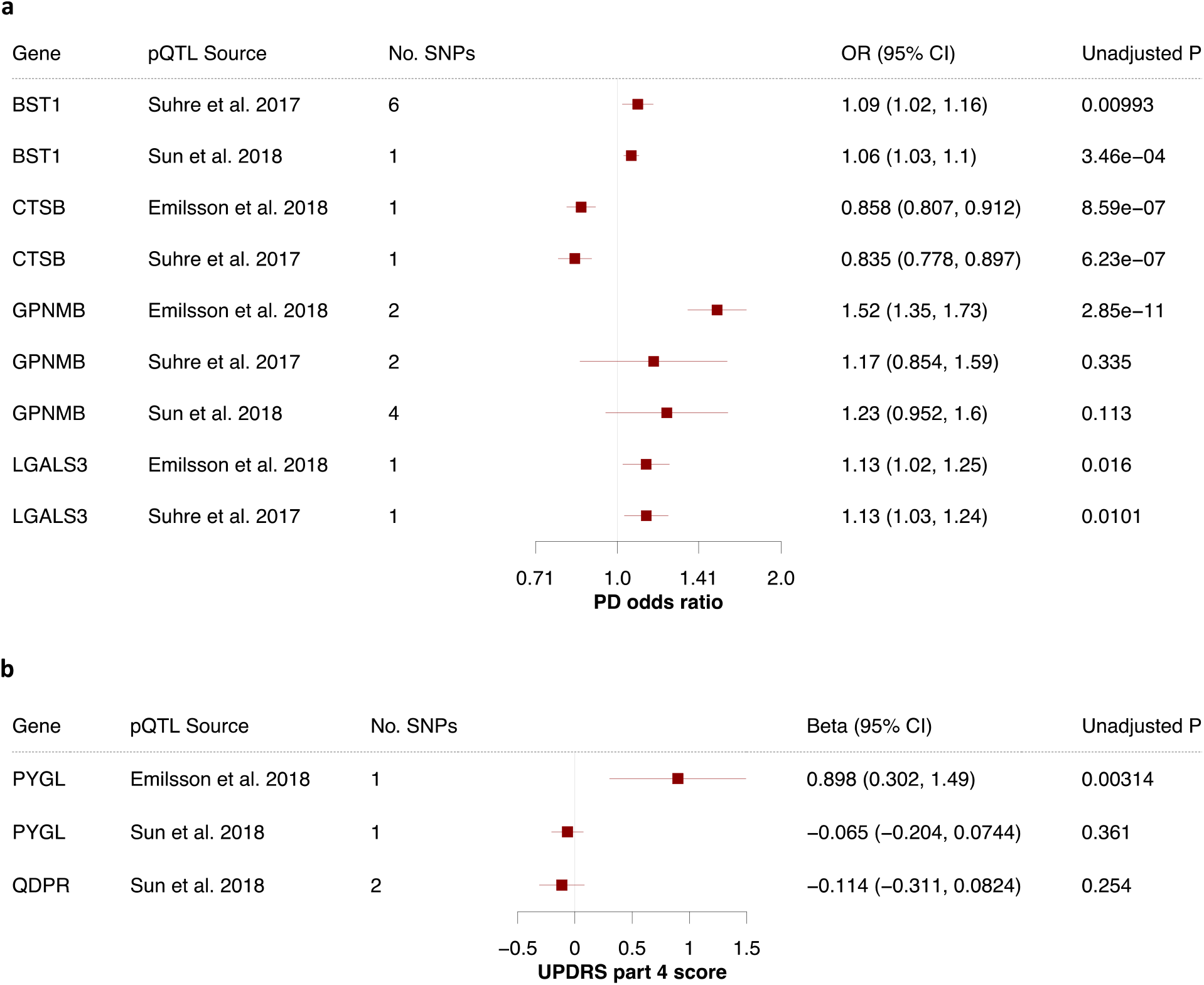
Protein quantitative trait loci in blood provide further genetic evidence. Forest plots showing the results for all proteins and outcomes where a pQTL was available. Results from the Wald ratio or IVW are shown for a) PD risk and b) UPDRS part 4. All pQTLs were measured in blood, and the “pQTL Source” column indicates which pQTL study the SNPs were derived from. 95% CI, 95% confidence interval; OR, odds ratio; pQTL, protein quantitative trait locus; UPDRS, unified PD rating scale.

It is noteworthy that *expression* of *CTSB, GPNMB* and *LGALS3* in *brain tissue* was significantly associated with PD risk, and here we find that levels of the encoded proteins in *blood* may also predict PD risk. The direction of effect was consistent between the pQTL and eQTL results for all genes except BST1. We found that BST1 protein levels in blood were associated with a raised PD risk, whereas raised expression of *BST1* was associated with reduced PD risk. Using the expression data, *BST1* did not pass the MR-Egger intercept test nor the *I*^2^ test. Similarly, when using the pQTL data provided by Suhre and colleagues for BST1, the MR-Egger intercept and Cochran’s Q test suggest that this result may be biased by genetic pleiotropy. This illustrates the importance of MR quality control tests for heterogeneity and directional pleiotropy, which is not possible when there is only one SNP available. Maximizing the number of SNPs available per drug target and validating drug targets with different data types and independent replication cohorts is essential.

## Discussion

### Prioritizing drug targets based on the strength of evidence

To our knowledge, this is the first MR study to date explicitly seeking to identify new drug targets for PD, and we provide genetic evidence for 23 potential disease-modifying drug targets. Of these, we consider those that pass MR quality control to have the most robust MR evidence. For PD risk, these are encoded by *CD38, GPNMB* and *MAP3K12*. For PD progression markers, we found the strongest MR evidence for *IRAK3* (dyskinesia), *LMAN1* (UPDRS part 2), *QDPR* (UPDRS part 4) and *RHD* (dyskinesia).

Raised *CD38* expression was associated both with a reduced PD risk and a delayed age at onset, and we find evidence that GPNMB protein levels in blood may significantly influence PD risk. *CD38* and *GPNMB* are furthermore supported by existing colocalization evidence, showing that the eQTLs for *CD38* and *GPNMB* in brain tissue colocalize with PD risk loci (Kia et al. 2020). There is evidence using protein QTL data that a protein with both MR and colocalization evidence is more likely to be a successful drug target (Zheng et al. 2019), and so *CD38* and *GPNMB* may be the most promising drug target candidates for PD risk.

Similarly, *QDPR* (UPDRS part 4) is further supported by pQTL data. We found that raised *IRAK3* expression may be protective against the development of dyskinesias, depression and progression of UPDRS parts 2 and 4 in PD. Likewise, raised *LMAN1* also predicted a lower UPDRS part 2 score. This strengthens the evidence for QDPR, IRAK3 and LMAN1 as potential targets of disease-modifying drugs.

We identify two drug classes which may increase PD risk: CD38 inhibitors and MAP3K12 inhibitors. CD38-inhibitors such as daratumumab, licensed to treat multiple myeloma, may increase PD risk and promote and earlier age at onset. We also find that inhibitors of the MAP3K12 protein may increase PD risk. Notably, the MAP3K12 inhibitor CEP-1347 showed encouraging evidence from animal studies as a medication to *treat* PD, and this drug failed to modify PD progression in a phase 3 clinical trial (Parkinson Study Group PRECEPT Investigators 2008). As such, our data provide a genetic explanation why CEP-1347 was unsuccessful.

### Four drug repurposing candidates

We identify four candidates for drug repurposing, of which one may modify PD risk and one may mitigate the development of dyskinesias (Table 1). These drugs could reach patients sooner and at a reduced cost, because they have already passed safety testing in humans. To our knowledge, there is no evidence linking PD and the drug roledumab, which is currently in a phase II clinical trial to prevent alloimmunisation in Rhesus negative mothers carrying a Rhesus positive child (NCT02287896). *NDUFAF2* on the other hand encodes a subunit of a target of metformin, an approved medication for type 2 diabetes mellitus, and there is extensive evidence for a relationship between diabetes and PD (Foltynie and Athauda 2020). Epidemiological studies studying the relationship between long-term medication use and incidence of a disease are an invaluable contribution to evaluating preventative agents for PD. A retrospective cohort study of over 6000 patients with type 2 diabetes mellitus found that more than four years of metformin use is associated with a reduced PD incidence (Shi et al. 2019). Together with this MR study, this provides further evidence in favour of repurposing anti-diabetic drugs for PD.

Other medications may not be as suitable for repurposing. *CHRNB1* encodes the beta subunit of the muscle acetylcholine receptor at the neuromuscular junction, which is inhibited by muscle relaxants used during surgical anaesthesia. *VKORC1* encodes the catalytic subunit of the vitamin K epoxide reductase, and this enzyme is targeted by the oral anticoagulant warfarin. The key adverse effect of warfarin treatment is haemorrhage, and since PD is a movement disorder where patients experience frequent falls, any potential benefit of warfarin treatment would likely be outweighed by the added risk of haemorrhagic strokes and complications of bleeding.

### Insights from using different PD traits as the outcome

The two-sample MR design allows us to explore different tissues and PD traits, providing valuable information about drug target sites and potential outcomes. We propose different candidates to (1) prevent PD, (2) delay PD onset, and (3) slow PD progression (Figures 2, 3, 4). The preventative list carries the most robust evidence, because each gene reached significance in two large, independent cohorts. Replication is critical to validating scientific findings and eliminating false positives, and this has been an crucial lesson for genetic research (Hirshhorn et al. 2002, Chanock et al. 2007; Marigorta et al. 2018). Replication is not common practice in MR yet (Burgess, Foley, and Zuber 2018), and it is a key strength of our study. Although including all samples available in one analysis would maximise statistical power (Chanock et al. 2007; Huffman 2018), including independent discovery and replication cohorts allowed us to robustly validate our proposed drug targets. Indeed, since our overarching intention is to provide genetic evidence to improve success rates in clinical trials, we made this decision in order to minimise the number of false positives.

Nevertheless, PD cannot be accurately predicted yet, and a preventative agent would need to be highly tolerable and have a very safe side effect profile. For these reasons, the list of candidates to slow PD progression may be very useful, despite the added challenge that measuring these outcomes is more subjective. Indeed, clinical trials generally use a progression marker as the primary outcome. Whereas it is more clear how the list of drug targets to slow PD progression are clinically relevant, the preventative drug targets we propose carry more robust evidence.

Moreover, many GWAS loci nominated to affect PD risk are not associated with age of onset or progression markers, painting the picture of different underlying molecular mechanisms (Nalls et al. 2019; Blauwendraat et al. 2019; Iwaki et al. 2019). On the one hand, we find that four of the drug targets that replicated for PD risk may also affect PD age at onset, suggesting a shared molecular pathophysiology. On the other hand, there is little overlap between our candidates for PD risk and progression, suggesting that different mechanisms may drive PD incidence versus clinical progression. As such, our data warrant further investigation into the relationship between the molecular mechanisms of PD susceptibility and progression.

Notwithstanding, the majority of our candidates were not proposed by any of the latest GWASs for PD traits (Nalls et al. 2019; Blauwendraat et al. 2019; Iwaki et al. 2019). This is expected, because GWAS identifies SNPs associated with a trait, and usually nominates genes close to the SNP or using an outcome-centred MR approach. For example, Nalls and colleagues selected SNPs associated with the outcome (PD risk) and used MR to identify whether any of these loci act through altering expression or methylation of genes within 1 Mb of the SNP. This contrasts with our exposure-centred MR analysis, where we chose SNPs associated with the exposure (druggable gene expression or protein levels). Our SNPs do not need to be close to a locus that is strongly associated with a PD trait. In fact, we have removed any SNPs that are more strongly associated with the PD trait than gene expression or protein levels from our study.

### How well do different QTL data mimic medications?

The choice of SNPs furthermore dictates how accurately this study mimics medications. We have restricted our eQTL analysis to SNPs within 5 kb of the associated gene to reduce the heterogeneity between SNPs. It is believed that eQTLs acting in *cis* (e.g. found within 1 Mb) of the linked gene are less pleiotropic than eQTLs acting in *trans* (Schmidt et al. 2020). Clumping at a very conservative threshold (e.g. *r*^2^ < 0.001) often leaves one or two SNPs (Schmidt et al. 2020), and this makes it difficult to test for pleiotropy and may yield type I errors (false positives). Nevertheless, some MR meta-analysis methods require strictly independent SNPs (e.g. mode-based and median-based methods), so a liberal clumping threshold makes it more difficult to probe inconsistencies between methods. We therefore clumped at *r*^2^ = 0.2 for our initial screen, and repeated the analysis for significant genes clumping at *r*^2^ = 0.001 (Burgess et al. 2019). This maximised our ability to test for heterogeneity between SNPs and directional pleiotropy in our main analysis, and we scrutinised our findings using a stricter clumping threshold.

The eQTL cohorts contain some non-European individuals (Võsa et al. 2018, Wang et al. 2018), whereas the PD GWAS populations are comprised of European individuals only (Nalls et al. 2019; Blauwendraat et al. 2019; Iwaki et al. 2019). Two of the pQTL studies soourced were based on Icelandic and German cohorts (Emilsson et al. 2018; Suhre et al. 2017). Linkage disequilibrium patterns differ between populations, and this may compromise how well our QTLs mimic drug action in the PD GWAS cohorts and introduce bias to the MR effect estimate (Burgess et al. 2019).

Another key limitation is that this MR study does not fully recapitulate a clinical trial. MR mimics lifelong, low-dose exposure to a drug and assumes a linear relationship between exposure and outcome. This differs from a clinical trial, which typically investigates comparably high doses a drug over a much shorter timeframe. The MR result may therefore not directly correspond to the effect size in practice and does not perfectly predict the effect of a drug.

Most medications target proteins, and it is unclear whether gene expression adequately mimics such drug action. eQTL datasets generally have large sample sizes, probe many genes and cover diverse tissues, and are ideal for a high-throughput screen. This is not the case for protein QTL datasets, limiting the possibilities to conduct a thorough MR study using protein data. We are encouraged that three of the six proteins we were able to probe using pQTL data were successfully validated. This study will add to existing evidence that regulatory variants may be used for robust causal inferences in drug target MR (Schmidt et al. 2020). Nevertheless, neither QTL type reflects activity levels of the protein, and this MR study does not provide functional evidence for the proposed drug targets.

It is difficult to interpret which tissue would be the most appropriate site of action. Whereas the genes which reached significance in both blood and brain tissue may have stronger MR evidence, targeting the protein of a widely expressed gene may lead to systemic side-effects. Brain tissue may be more biologically relevant for neurodegeneration, but a drug acting in the blood stream may not need to cross the blood brain barrier to exert its effect. We have included both blood and brain tissue in order to capture as many genes as possible and explore two potential tissue sites of action, but it is difficult to prioritise genes based on which tissue(s) they reached significance in.

### Conclusion

There is evidence that a 9.6% vs. 13.8% success rate for drugs from phase 1 trials to approval may mean a $480 million difference in the median research and development cost required to bring a new drug to the market (Wouters, McKee, and Luyten 2020). Therefore, any genetic evidence which increases success rates even by a few percent may have a substantial effect on drug development costs (Nelson et al. 2015; King, Wade Davis, and Degner 2019). As such, MR is a highly compelling, time- and cost-effective adjunct to the randomized controlled trial, made possible by large-scale GWAS data. We make our code openly available for use beyond PD research (https://github.com/catherinestorm/mr_druggable_genome_pd/), and we demonstrate ways to prioritise drug targets based on genetic data. We provide human genetic evidence of drug efficacy for PD, and we hope that these data will serve as a useful resource for prioritising drug development efforts.

## Acknowledgements

CSS would like to thank Dr Vishal Rawji for his invaluable support and insightful ideas about the clinical implications and communication of this study. CSS is funded by Rosetrees Trust, John Black Charitable Foundation and the University College London MBPhD Programme. DAK is supported by an MBPhD Award from the International Journal of Experimental Pathology. MA is funded by the Faculty of Applied Medical Sciences, King Abdulaziz University, Jeddah, Saudi Arabia. NWW is a National Institute for Health Research senior investigator and receives support from the European Union Joint Programme—Neurodegenerative Disease Research Medical Research Council Comprehensive Unbiased Risk factor Assessment for Genetics and Environment in Parkinson’s disease. ADH is an NIHR Senior Investigator. NWW, ADH and CF receive support from the National Institute for Health Research University College London Hospitals Biomedical Research Centre. We would like to thank all members of the International Parkinson Disease Genomics Consortium (IPDGC) and the authors of QTL projects referenced here, who make their data openly available. We thank all the patients and families whose decision to donate tissue samples make our research possible.

## Author Contributions

Conceptualization, CSS, DAK, MA, NWW; Methodology, CSS, DAK, MA, NWW; Investigation, formal analysis, visualization, CSS; Resources, SBC, CF, ADH, IPDGC; Writing – Original Draft, CSS; Writing – Review & Editing, CSS, DAK, MA, NWW, SBC, CF, ADH, IPDGC.

## Declaration of Interests

The authors report no competing interests.

## Methods

All DNA positions are based on the human reference genome build 37. Data processing was completed using R software version 3.6.3 (R Core Team 2019).

### Exposure data

Tissue-specific eQTL data were obtained from the eQTLGen (https://eqtlgen.org/) and PsychENCODE consortia (http://resource.psychencode.org/); full descriptions of the data are available in the original publications (Võsa et al. 2018; Wang et al. 2018). Briefly, the eQTLGen data consist of cis-eQTLs for 16,987 genes and 31,684 blood samples, of which most are healthy European-ancestry individuals. We downloaded the full significant cis-eQTL results (FDR < 0.05) and allele frequency information from the eQTLGen consortium on May 13th 2020.

The PsychENCODE data include 1,387 prefrontal cortex primarily-European samples (679 healthy controls, 497 schizophrenia, 172 bipolar disorder, 31 autism spectrum disorder and 8 affective disorder patients). We downloaded all significant eQTLs (FDR < 0.05) for genes with expression > 0.1 fragments per kilobase per million mapped fragments (FPKM) in at least 10 samples and all SNP information, accessed on May 13th 2020.

We obtained an updated version of the druggable genome containing 4,863 genes through personal correspondence with the authors of the original publication (Finan et al. 2017), double-checking the druggability level for all genes marked as approved or in clinical trials (“druggability tier 1”). We removed non-autosomal genes, leaving 4,560 druggable genes. We filtered both eQTL datasets to include SNPs 5 kb upstream of the target druggable gene start or 5 kb downstream of the target druggable gene end position.

We sought freely available pQTL data from blood or brain tissue for all druggable genes which reached significance for any PD outcome in our study. Out of 23 pQTL studies identified, three studies (1) reported significant pQTLs in individuals of European descent for any of the druggable proteins proposed by our eQTL analysis, (2) provided all the SNP information required for MR and (3) reports SNPs that were available in our PD outcome data.

Sun and colleagues measured 3,622 proteins in 3,301 healthy European blood donors from the INTERVAL study and identified 1,927 pQTLs for 1,478 protein. Emilsson and colleagues measured 4,137 proteins in the serum of 5,457 Icelanders from AGES Reykjavik study. Effect alleles and effect allele frequencies were obtained through personal correspondence with the authors. Suhre and colleagues measured 1,124 proteins in 1,000 blood samples from a German population.

In total, we found pQTLs that were available in the appropriate PD outcome data for six of our druggable proteins of interest: BST1, CTSB, GPNMB, LGALS3, PYGL, and QDPR. All pQTLs included in our analysis had *p* < 5e-6 in the original pQTL study. All pQTLs were found on the same chromosome as the associated gene except for: rs62143198 for PYGL, rs62143197 for QDPR, rs4253282 for GPNMB, rs2731674 for GPNMB (Sun et al. 2018).

### Outcome data

All PD data were obtained from the IPDGC, and details on recruitment and quality control are available in the original publications. In the discovery phase for PD risk we used openly available summary statistics from a 2014 case-control GWAS meta-analysis, which includes 13,708 PD patients and 95,282 controls (Nalls et al. 2014).

In the replication phase for PD risk, we obtained summary statistics from 11 case-control GWAS studies included in the most recent PD risk GWAS meta-analysis from the authors (Nalls et al. 2019). The 11 studies, as named and described in the PD GWAS meta-analysis, were Spanish Parkinson’s, Baylor College of Medicine/University of Maryland, McGill Parkinson’s, Oslo Parkinson’s Disease Study, Parkinson’s Progression Markers Initiative (PPMI), Finnish Parkinson’s, Harvard Biomarker Study (HBS), UK PDMED (CouragePD), Parkinson’s Disease Biomarker’s Program (PDBP), Tubingen Parkinson’s Disease cohort (CouragePD) and Vance (dbGap phs000394). These yielded a total of 8,036 PD cases and 5,803 controls. We meta-analysed the data using METAL (version 2011-03-25) using default settings, weighted by sample size (Willer, Li, and Abecasis 2010). The overall genomic inflation factor was *λ* = 1.116, and when scaled to 1,000 cases and 1,000 controls λ_1000_ = 1.017. The quantile-quantile plot showed adequate agreement with the expected null distribution (Figure S1).

For the progression marker analyses, we used summary statistics from the largest publicly available GWAS meta-analyses for PD age at onset and clinical progression (Blauwendraat et al. 2019; Iwaki et al. 2019). For age at onset, this includes 17,996 PD cases, and age at onset was defined as self-reported age at motor symptom onset or PD diagnosis. The authors reported a high correlation between age of diagnosis and age at onset.

The progression GWAS meta-analysis included 4,093 PD patients from 12 cohorts, followed over a median of 2.97 years (mean visits per individual over the study period: 5.44). We downloaded summary statistics for nine continuous outcomes and four binomial outcomes (). Continuous outcomes included Hoehn and Yahr stage (PD progression rating scale), total UPDRS/Movement Disorder Society revised version total (PD progression rating scale), UPDRS parts 1 to 4 (1 = non-motor symptoms, 2 = motor symptoms, 3 = motor examination, 4 = motor complications), MOCA (cognitive impairment), MMSE (cognitive impairment), SEADL (activities of daily living and independence). The binomial outcomes we used were: dementia, depression, dyskinesia, reading Hoehn and Yahr stage 3 or more.

### Mendelian randomization

MR analyses were completed using the R package “TwoSampleMR” version 0.5.4 (Hemani et al. 2018), unless stated otherwise. The exposure and outcome data were loaded and harmonized using in-built functions. SNPs were then clumped at *r*^2^ < 0.2 using European samples from the 1000 Genomes Project (Hemani et al. 2018; The 1000 Genomes Project Consortium 2012). Steiger filtering was used to remove SNPs that explain a greater proportion of variation in the outcome (PD trait) than variation in the exposure (gene expression).

Wald ratios were calculated for all SNPs. These were meta-analysed using the IVW, MR-Egger and maximum likelihood methods, including a linkage disequilibrium matrix to account for some correlation between SNPs; this function uses the R package “MendelianRandomization” version 0.4.2 (Yavorska and Burgess 2017). Forest plots were produced using the R package “forestplot”.

Where > 2 SNPs were available per exposure, we used the MR-Egger method and assessed whether the MR-Egger intercept significantly deviated from zero, as well as Cochran’s Q and *I*^2^ tests for heterogeneity between Wald ratios. The *I*^2^ was calculated as shown below, where *Q* is Cochran’s Q and *n* is the number of Wald ratios meta-analysed.

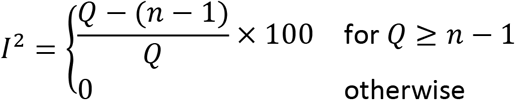

For genes which reached significance using the IVW method (> 1 SNP available), we carried out another MR analysis, clumping at *r*^2^ < 0.001. If > 1-2 SNPs were available at this clumping threshold, Wald ratios were meta-analysed using the IVW, MR-Egger, weighted mode and weighted median methods.

## Resource Availability

### Lead Contact

Further information and requests for resources and data should be directed to and will be fulfilled by the Lead Contact, Nicholas W Wood.

### Materials Availability

Not applicable.

### Data and Code availability

The code generated for this study is openly available on GitHub, accompanied by instructions for use (https://github.com/catherinestorm/mr_druggable_genome_pd). The supplementary information contains full results, and interim data can be requested by contacting the corresponding authors. The openly available data used in this study can be accessed as described in the methods section, and access to all other data used in this study is regulated by the authors of the original publications, referenced in the methods section.

## Supplementary Information

Table S1. Related to Figures 2, 3 and 4. MR results for all genes reaching significance for PD risk (discovery phase and replication phase), age at onset and progression markers.

Table S2. Related to Figures 2, 3 and 4. MR quality control (MR Egger intercept, Cochran’s Q, *I*^2^ tests) results for all genes reaching significance for PD risk (discovery phase and replication phase), age at onset and progression markers.

Table S3. Related to Figures 2, 3 and 4. MR results for all genes tested for PD risk (discovery phase and replication phase), age at onset and progression markers.

Table S4. Related to Figures 2, 3 and 4. MR quality control results (MR Egger intercept, Cochran’s Q, *I*^2^ tests) for all genes tested for PD risk (discovery phase and replication phase), age at onset and progression markers.

Table S5. Related to Figure 5. MR results for all proteins where a pQTL was available.

Table S6. Related to Figure 5. R quality control results (MR Egger intercept, Cochran’s Q, *I*^2^ tests) for all proteins where a pQTL was available.

Figure S1. Quantile-quantile plot for the meta-analysis of PD case-control GWAS data used for the replication step for PD risk.

## Notes

### Competing Interest Statement

The authors have declared no competing interest.

## References

Blauwendraat, Cornelis, Karl Heilbron, Costanza L Vallerga, Sara Bandres-ciga, Rainer Von Coelln, Lasse Pihlstrøm, Javier Simón-sánchez, et al. 2019. “Parkinson’s Disease Age at Onset Genome-Wide Association Study: Defining Heritability, Genetic Loci, and α-Synuclein Mechanisms.” Mov. Disord., 1–10. https://doi.org/10.1002/mds.27659.

Burgess, Stephen, George Davey Smith, Neil M. Davies, Frank Dudbridge, Dipender Gill, M. Maria Glymour, Fernando P. Hartwig, et al. 2019. “Guidelines for performing Mendelian randomization investigations.” Wellcome Open Res. 4: 186. https://doi.org/10.12688/wellcomeopenres.15555.1.

Burgess, Stephen, Frank Dudbridge, and Simon G. Thompson. 2016. “Combining information on multiple instrumental variables in Mendelian randomization: Comparison of allele score and summarized data methods.” Stat. Med. 35 (11): 1880–1906. https://doi.org/10.1002/sim.6835.

Burgess, Stephen, Christopher N Foley, and Verena Zuber. 2018. “Inferring Causal Relationships Between Risk Factors and Outcomes from Genome-Wide Association Study Data Stephen.” Annu. Rev. Genom. Hum. Genet., no. 19: 303–27. https://doi.org/10.1146/annurev-genom-083117-021731-021731.

Chanock, Stephen J., Teri Manolio, Michael Boehnke, Eric Boerwinkle, David J. Hunter, Gilles Thomas, Joel N. Hirschhorn, et al. 2007. “Replicating genotype-phenotype associations.” https://doi.org/10.1038/447655a.

Emilsson V, Ilkov M, Lamb JR, Finkel N, Gudmundsson EF, Pitts R, et al. 2018 “Co-regulatory networks of human serum proteins link genetics to disease.” Science. 361: 769–773.

Escott-Price, Valentina, Mike A. Nalls, Huw R. Morris, Steven Lubbe, Alexis Brice, Thomas Gasser, Peter Heutink, et al. 2015. “Polygenic risk of Parkinson disease is correlated with disease age at onset.” Ann. Neurol. 77 (4): 582–91. https://doi.org/10.1002/ana.24335.

Finan, Chris, Anna Gaulton, Felix A Kruger, R Thomas Lumbers, Tina Shah, Jorgen Engmann, Luana Galver, et al. 2017. “The druggable genome and support for target identification and validation in drug development” 1166 (March): 1–15.

Foltynie T, Athauda D. 2020. “Repurposing anti-diabetic drugs for the treatment of Parkinson’s disease: Rationale and clinical experience.” Progress in Brain Research. Prog Brain Res 252:493–523. https://doi.org/10.1016/bs.pbr.2019.10.008

Giambartolomei C. et al. 2014. “Bayesian test for colocalisation between pairs of genetic association studies using summary statistics.” PLoS Genet., 10, e1004383.

The GTEx Consortium. 2015 “The Genotype-Tissue Expression (GTEx) pilot analysis: Multitissue gene regulation in humans.” Science. 348(6235): 648–60.

Harrison, Richard K. 2016. “Phase II and phase III failures: 2013 – 2015.” Nat. Rev. Drug Discov. 15 (12): 817–18. https://doi.org/10.1038/nrd.2016.184.

Haycock, Philip C, Stephen Burgess, Kaitlin H Wade, Jack Bowden, Caroline Relton, and George Davey Smith. 2016. “Best (but oft-forgotten) practices: the design, analysis, and interpretation of Mendelian randomization studies.” Am J Clin Nutr 103 (February): 965–78. https://doi.org/10.3945/ajcn.115.118216.INTRODUCTION.

Hemani, Gibran, Jie Zheng, Benjamin Elsworth, Kaitlin H Wade, Valeriia Haberland, Denis Baird, Charles Laurin, et al. 2018. “The MR-Base platform supports systematic causal inference across the human phenome.” Elife 7: e34408. https://doi.org/10.7554/eLife.34408.

Hemani G, Bowden J, Davey Smith G. 2018a. Evaluating the potential role of pleiotropy in Mendelian randomization studies. Hum Mol Genet. 27: R195–R208.

Hingorani AD, Humphries SE. “Nature’s randomised trials.” 2005. Lancet. 366: 1906–1908.

Hingorani AD, Kuan V, Finan C, Kruger FA, Gaulton A, Chopade S, et al. 2019. “Improving the odds of drug development success through human genomics: modelling study.” Sci Rep. 9: 1–25.

Hirschhorn JN, Lohmueller K, Byrne E, Hirschhorn K. 2002 “A comprehensive review of genetic association studies.” Genet Med. 4: 45–61.

Holmes M V., Ala-Korpela M, Davey Smith G. 2017. “Mendelian randomization in cardiometabolic disease: challenges in evaluating causality.” Nat Rev Cardiol. 14: 577–590.

Huffman JE. 2018. “Examining the current standards for genetic discovery and replication in the era of mega-biobanks.” Nat Commun. 9: 1–4.

Ibanez, Laura, Umber Dube, Benjamin Saef, John Budde, Kathleen Black, Alexandra Medvedeva, Jorge L. Del-Aguila, et al. 2017. “Parkinson disease polygenic risk score is associated with Parkinson disease status and age at onset but not with alpha-synuclein cerebrospinal fluid levels.” BMC Neurol. 17 (1): 1–9. https://doi.org/10.1186/s12883-017-0978-z.

Iwaki, Hirotaka, Cornelis Blauwendraat, Hampton L. Leonard, Ganqiang Liu, Jodi Maple-Grødem, Jean Christophe Corvol, Lasse Pihlstrøm, et al. 2019. “Genetic risk of Parkinson disease and progression: An analysis of 13 longitudinal cohorts.” Neurol. Genet. 5 (4). https://doi.org/10.1212/NXG.0000000000000348.

Katan, Martijn B. 1986. “Apoliporotein E isoforms, serum cholesterol, and cancer.” Lancet, no. March: 507–8.

Kia DA, Zhang D, Guelfi S, Manzoni C, Hubbard L, UKBEC, et al. 2020. “Integration of eQTL and Parkinson’s disease GWAS data implicates 11 disease genes.” JAMA Neurol. Accepted.

King, Emily A., J. Wade Davis, and Jacob F. Degner. 2019. “Are drug targets with genetic support twice as likely to be approved? Revised estimates of the impact of genetic support for drug mechanisms on the probability of drug approval.” PLoS Genet. 15 (12): 1–20. https://doi.org/10.1371/journal.pgen.1008489.

Marigorta, Urko M., Juan Antonio Rodríguez, Greg Gibson, and Arcadi Navarro. 2018. “Replicability and Prediction: Lessons and Challenges from GWAS.” Trends Genet. 34 (7): 504–17. https://doi.org/10.1016/j.tig.2018.03.005.

Nalls, Mike A., Cornelis Blauwendraat, Costanza L. Vallerga, Karl Heilbron, Sara Bandres-Ciga, Diana Chang, Manuela Tan, et al. 2019. “Identification of novel risk loci, causal insights, and heritable risk for Parkinson’s disease: a meta-analysis of genome-wide association studies.” Lancet Neurol. 18 (12): 1091–1102. https://doi.org/10.1016/S1474-4422(19)30320-5.

Nalls, Mike A., Valentina Escott-Price, Nigel M. Williams, Steven Lubbe, Margaux F. Keller, Huw R. Morris, and Andrew B. Singleton. 2015. “Genetic risk and age in Parkinson’s disease: Continuum not stratum.” Mov. Disord. 30 (6): 850–54. https://doi.org/10.1002/mds.26192.

Nalls, Mike A, Nathan Pankratz, Christina M Lill, Chuong B Do, Dena G Hernandez, Mohamad Saad, Anita L DeStefano, et al. 2014. “Large-scale meta-analysis of genome-wide association data identifies six new risk loci for Parkinson’s disease.” Nat. Genet. 56 (9): 1–7. https://doi.org/10.1038/ng.3043.

Nelson, Matthew R, Hannah Tipney, Jeffery L Painter, Judong Shen, Paola Nicoletti, Yufeng Shen, Aris Floratos, et al. 2015. “The support of human genetic evidence for approved drug indications.” Nat. Publ. Gr. 47 (8): 856–60. https://doi.org/http://dx.doi.org/10.1038/ng.3314.

R Core Team. 2019. “R: A Language and Environment for Statistical Computing.” R Foundation for Statistical Computing, Vienna, Austria. http://www.R-project.org/

Ramasamy A, Trabzuni D, Guelfi S, Varghese V, Smith C, Walker R, et al. 2014. “Genetic variability in the regulation of gene expression in ten regions of the human brain.” Nat Neurosci. 17(10): 1418–28.

Rose AAN, Biondini M, Curiel R, Siegel PM. 2017. “Targeting GPNMB with glembatumumab vedotin: Current developments and future opportunities for the treatment of cancer.” Pharmacol Ther. 179: 127–141.

Parkinson Study Group PRECEPT Investigators. 2008. “Mixed lineage kinase inhibitor CEP-1347 fails to delay disability in early parkinson disease.” Neurology 71 (6): 462–63. https://doi.org/10.1212/01.wnl.0000324506.93877.5e.

Schmidt AF, Finan C, Gordillo-Marañón M, Asselbergs FW, Freitag DF, Patel RS, et al. 2020. “Genetic drug target validation using Mendelian randomisation.” Nat Commun.

Shi Q, Liu S, Fonseca VA, Thethi TK, Shi L. 2019. “Effect of metformin on neurodegenerative disease among elderly adult US veterans with type 2 diabetes mellitus.” BMJ Open.

Slob, Eric A. W., and Stephen Burgess. 2020. “A comparison of robust Mendelian randomization methods using summary data.” Genet. Epidemiol., no. April 2019: 1–17. https://doi.org/10.1002/gepi.22295.

Smietana, Katarzyna, Marcin Siatkowski, and Martin Møller. 2016. “Trends in clinical success rates.” Nat. Rev. Drug Discov. 15 (6): 379–80. https://doi.org/10.1038/nrd.2016.85.

Smith, George Davey, and Shah Ebrahim. 2003. “‘Mendelian randomization’: can genetic epidemiology contribute to understanding environmental determinants of disease?*.” Int. J. Epidemiol. 32: 1–22. https://doi.org/10.1093/ije/dyg070.

Storm CS, Kia DA, Almramhi M, Wood NW. 2020. “Using Mendelian randomization to understand and develop treatments for neurodegenerative disease.” Brain Commun.

Suhre K, Arnold M, Bhagwat AM, Cotton RJ, Engelke R, Raffler J, et al. 2017 “Connecting genetic risk to disease end points through the human blood plasma proteome.” Nat Commu. 8

Sun BB, Maranville JC, Peters JE, Stacey D, Staley JR, Blackshaw J, et al. 2018. “Genomic atlas of the human plasma proteome.” Nature. 558: 73–79.

The 1000 Genomes Project Consortium. 2012. “An integrated map of genetic variation from 1,092 human genomes.” Nature 491 (7422): 56–65. https://doi.org/10.1038/nature11632.

Võsa, Urmo, Annique Claringbould, Harm-Jan Westra, Marc Jan Bonder, Patrick Deelen, Biao Zeng, Holger Kirsten, et al. 2018. “Unraveling the polygenic architecture of complex traits using blood eQTL metaanalysis.” bioRxiv, 447367. https://doi.org/10.1101/447367.

Wang, Daifeng, Shuang Liu, Jonathan Warrell, Hyejung Won, Xu Shi, Fabio C. P. Navarro, Declan Clarke, et al. 2018. “Comprehensive functional genomic resource and integrative model for the human brain.” Science (80-.). 362 (6420). https://doi.org/10.1126/science.aat8464.

Willer, Cristen J., Yun Li, and Gonçalo R. Abecasis. 2010. “METAL: Fast and efficient meta-analysis of genomewide association scans.” Bioinformatics 26 (17): 2190–1. https://doi.org/10.1093/bioinformatics/btq340.

Wouters, Olivier J., Martin McKee, and Jeroen Luyten. 2020. “Estimated Research and Development Investment Needed to Bring a New Medicine to Market, 2009-2018.” JAMA - J. Am. Med. Assoc. 323 (9): 844–53. https://doi.org/10.1001/jama.2020.1166.

Yavorska, Olena O., and Stephen Burgess. 2017. “MendelianRandomization: An R package for performing Mendelian randomization analyses using summarized data.” Int. J. Epidemiol. 46 (6): 1734–9. https://doi.org/10.1093/ije/dyx034.

Zheng J, Haberland V, Baird D, Walker V, Haycock P, Richardson TG, et al. 2019. Phenome-wide Mendelian randomization mapping the influence of the plasma proteome on complex diseases. bioRxiv. https://doi.org/10.1101/627398

Zhu Z, Zhang F, Hu H, Bakshi A, Robinson MR, Powell JE, et al. 2016 Integration of summary data from GWAS and eQTL studies predicts complex trait gene targets. Nat Genet; 48: 481–487.

