## Supplementary figures and images for "Finding drug targeting mechanisms with genetic evidence for Parkinson’s disease"

### Figure S1

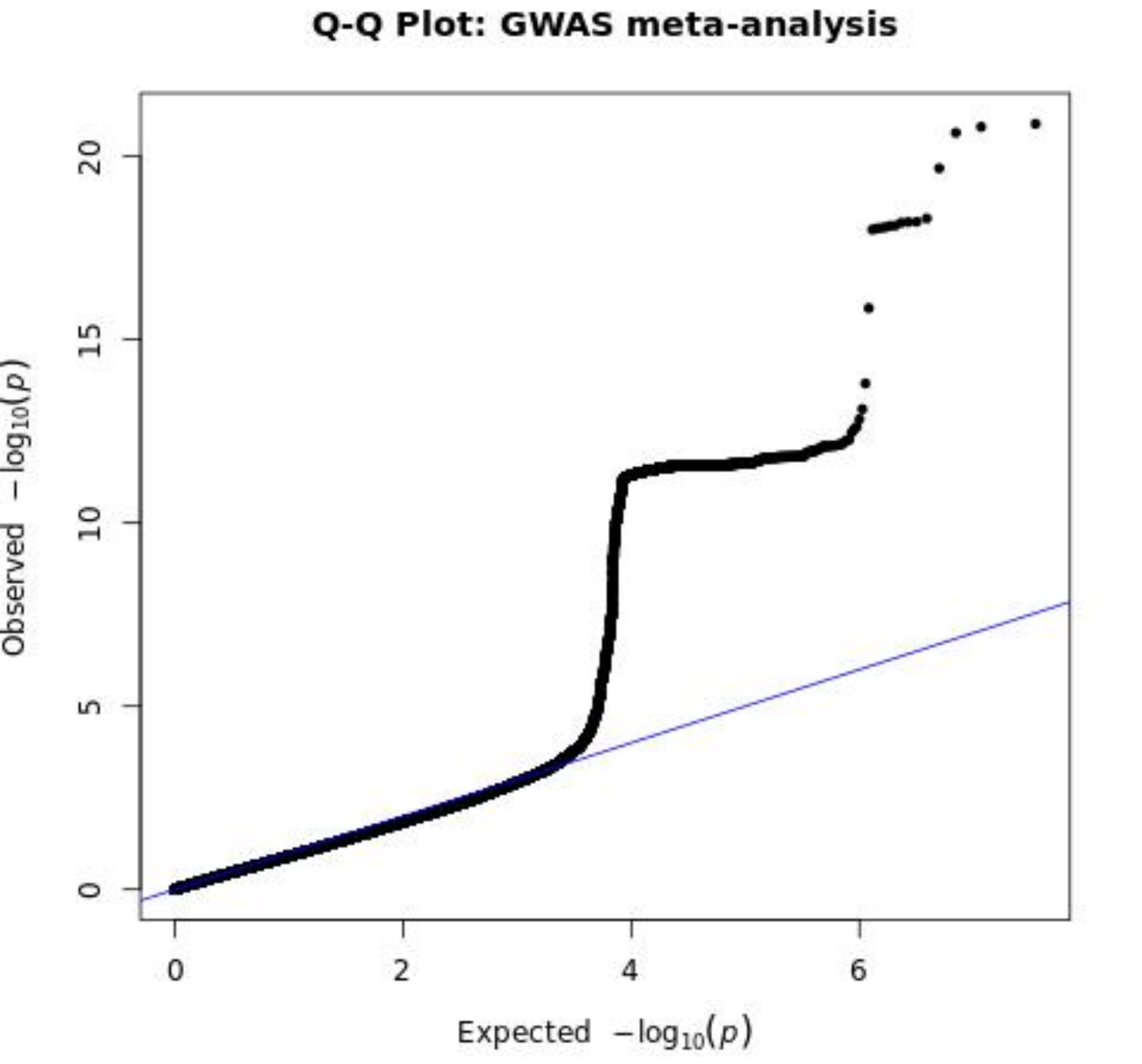
